# MCL1 dependence across MDS subtypes and dual inhibition of MCL1 and BCL2 in MISTRG6 mice

**DOI:** 10.1101/2020.06.05.133090

**Authors:** Melissa A. Fischer, Yuanbin Song, Rana Gbyli, Maria P. Arrate, Matthew T. Villaume, Merrida A. Childress, Brianna N. Smith, Thomas P. Stricker, Stephanie Halene, Michael R. Savona

## Abstract

Treatment for myelodysplastic syndromes (MDS) remains insufficient due to clonal heterogeneity and clinical complexity. Dysregulation of apoptosis is observed across MDS subtypes regardless of mutations and represents an attractive therapeutic opportunity. Venetoclax (VEN), a selective inhibitor of anti-apoptotic protein B-cell lymphoma-2 (BCL2), has yielded impressive responses in older patients with acute myeloid leukemia (AML). BCL2 family anti-apoptotic proteins BCL-X_L_ and induced myeloid cell leukemia 1 (MCL1) are implicated in leukemia survival, and upregulation of MCL1 is seen in VEN-resistant AML and MDS. Here, we determined the *in vitro* sensitivity of MDS patient samples to selective inhibitors of BCL2, BCL-X_L_ and MCL1. While VEN response positively correlated with increasing blast counts, all MDS subtypes responded to the MCL1 inhibitor, S63845. Treatment with combined VEN+S63845 was synergistic in all MDS subtypes and reduced MDS engraftment in MISTRG6 mice supporting the pursuit of clinical trials with combined BCL2+MCL1 inhibition in MDS.

## Introduction

Myelodysplastic syndromes (MDS) are heterogeneous bone marrow failure neoplasms marked by cytopenias, reduced quality of life and predilection to transform into acute myeloid leukemia (AML). Readily available treatments for MDS are lacking, and adapting newly approved therapies designed for AML to MDS is complicated, largely due to heterogeneity of MDS. Despite this, consistent with other myeloid malignancies, clonal hematopoietic stem and progenitor cells (HSPCs) in MDS avoid programmed cell death often share a pattern of an imbalance of mitochondrial-controlled BCL2 family proteins1-4. The BCL2 family includes anti-apoptotic and pro-apoptotic proteins that compete for ligand to block or promote the activation of BAX/BAK oligomerization that is required to induce mitochondrial outer membrane potential and subsequent apoptosis^5,6^. As a method to evade apoptosis, cancer cells often upregulate anti-apoptotic proteins BCL2 and MCL1, leading to a survival advantage in numerous malignancies^5-7^. Thus, selective targeting of anti-apoptotic proteins is a viable treatment strategy. Venetoclax (VEN), a newly FDA-approved therapy that specifically inhibits the anti-apoptotic protein, BCL2, has yielded response rates of up to 50-70% in elderly AML when combined with DNA methyltransferase inhibitors (DNMTi) or low dose cytarabine^8,9^. Upregulation of another BCL2 family anti-apoptotic protein, MCL1, is seen in AML and MDS treated with VEN, and is a noted mechanism of VEN-resistance.^4,10-12^ We previously revealed a selective MCL1 inhibitor with activity in AML patient samples dependent on MCL1 protein or resistant to BCL2 inhibition, including AML cells that arose from MDS.^11^ Taken together, targeting MDS cells via inhibition of BCL2 and/or MCL1 has appeal. We determined the sensitivity of MDS cells to inhibition of specific anti-apoptotic proteins, elucidated the characteristic determinants of response, and illustrated synergy with combined BCL2 and MCL1 inhibition both *in vitro* and *in vivo*.

## Results

### MDS subtypes are differentially sensitive to BCL2 inhibition and indiscriminately sensitive to MCL1 inhibition

We successfully cultured 35 MDS patient samples and determined their *in vitro* sensitivity to BCL2, BCL-X_L_ and MCL1 inhibitors. Using CellTiter-Glo, we determined the relative cell viability concentrations (GI_50_) for each inhibitor after 48 hours of exposure (Table 1). While few samples were sensitive to BCL-X_L_ inhibition (A-1155463), we observed a range of sensitivities to BCL2 inhibition that positively correlated with blast count. Consistent with previous findings,^1,4^ lower blast count MDS (RS-SLD/MLD and MLD) exhibited less sensitivity than higher blast count (EB1 and EB2) MDS subtypes (P = 0.01) (Supplementary Figure 1a). All subtypes were sensitive to the selective MCL1 inhibitor, S63845. Samples were assessed using a targeted next generation sequencing (NGS) panel of 37 commonly mutated genes in myeloid diseases. As expected, we observed an increased number of *SF3B1* mutations in the lower blast count MDS patient samples (MDS-RS)^13^. RAS-family mutants, particularly *PTPN11*-mutated AML have previously been shown to connote resistance to VEN^14-18^. In this small cohort, one of 2 *PTPN11* mutant MDS samples, and 2/2 *CBL* mutant samples were completely resistant to BCL2 inhibition (MDS033, MDS005 and MDS031, respectively). Otherwise, we did not observe any correlation between specific mutations and drug response in this cohort of 35 patient samples.

**Table 1.**
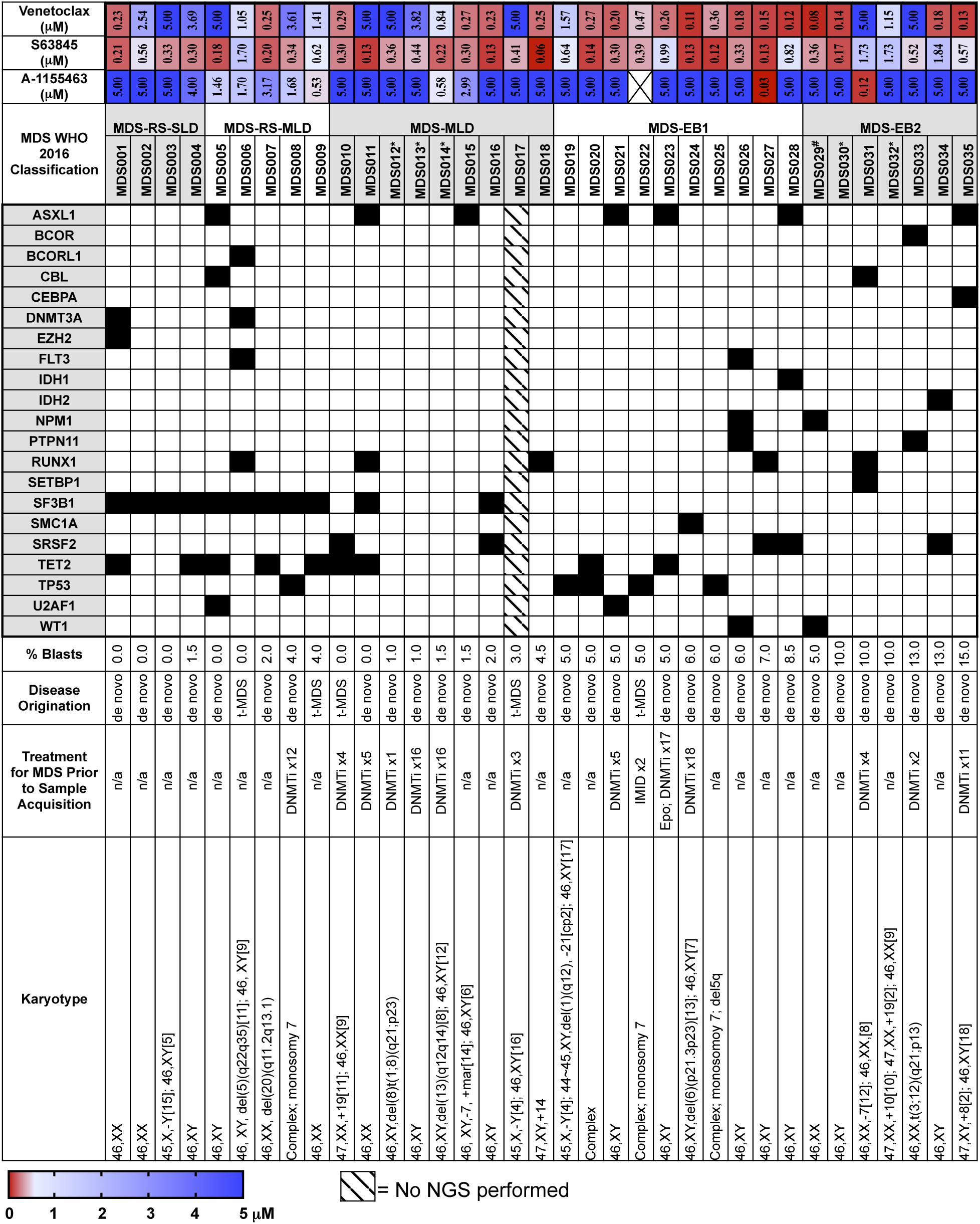
Comprehensive data for 21 untreated, and 14 previously treated MDS patient samples assessed is represented including growth inhibition at 50% (GI_50_) for each inhibitor, MDS subtype by 2016 WHO classification, sample name/number, mutational status, bone marrow blast percentage, disease origination, treatment for MDS prior to sample acquisition, and karyotype. * NGS mutation analysis was conducted but no mutation detected. #Patient MDS029 was considered MDS-EB2 despite only 5% blasts given the presence of Auer rods present in bone marrow aspirate. Other genes tested with the OnkoSight™ NGS Myeloid Malignancies gene panel that were not detected in any of the samples include ABL1, ATRX, BRAF, CALR, CBLB, CBLC, CDKN2A, CUX1, ETV6, FBXW7, GATA1, GATA2, GNAS, HRAS, IKZF1, JAK2, JAK3, KDM6A, KIT, KRAS, MLL, MPL, MYD88, NOTCH1, NRAS, PDGFRA, PHF6, PTEN, RAD21, SFS3R, SMC3, STAG2, ZRSR2.

### Dual inhibition of MCL1 and BCL2 is synergistic in all MDS samples resulting in increased apoptosis and loss of clonogenicity

To determine if co-inhibition of BCL2 and MCL1 is synergistic in MDS, we assessed the efficacy of three-fold dilutions of S63845+VEN on cell viability and employed the ZIP synergy model to determine the synergistic potential^19^. The combined treatment for all samples tested resulted in an average delta synergy score > 0, suggesting drug synergy (Fig. 1a and Supplementary Table 1). To investigate if the reduced viability seen with dual inhibition of BCL2 and MCL1 led to apoptosis of MDS CD34^+^ progenitor cells, cells were stained with anti-human CD34, Annexin V, and PI 24 hours after treatment with sub-therapeutic doses of each inhibitor (Supplementary Figure 1b). Cells treated with S63845+VEN displayed significant reductions in MDS progenitor cells compared to VEN-treated or S63845-treated cells when all MDS samples were analyzed together (Fig. 1b). Similarly, a refined analysis of MDS subtypes revealed that combination treatment significantly reduced MDS progenitors for all subtypes compared to cells treated with S63845 alone, or in RS-SLD/MLD and MLD for cells treated with VEN alone with a trend toward significance for EB1 and EB2 patient samples (Fig. 1c). To determine the long-term effects of reduced viability and increased apoptosis on clonogenicity, colony forming unit (CFU) assays were performed. In the tested patient samples, dual treatment reduced CFU-granulocyte magrophage (CFU-GM) formation by 50-82%, indicating loss of clonogenicity, even in a *CBL* (MDS031) or *PTPN11* (MDS033) mutant samples that were resistant to VEN monotherapy (82% and 50% reduction, respectively) (Fig. 1d). This was further confirmed by trypan blue staining on cells collected from the plates of each sample tested, which showed a reduction of total live cells (Supplementary Figure 1c). Given the skewing toward CFU-GM colonies from the MDS samples that had been cryopreserved (Fig. 1d), we also verified the effects of S63845+VEN treatment in freshly-obtained MDS patient samples, which corroborated a reduction, specifically, in CFU-GM colonies by 30-93% (Fig. 1e).

**Fig. 1.**
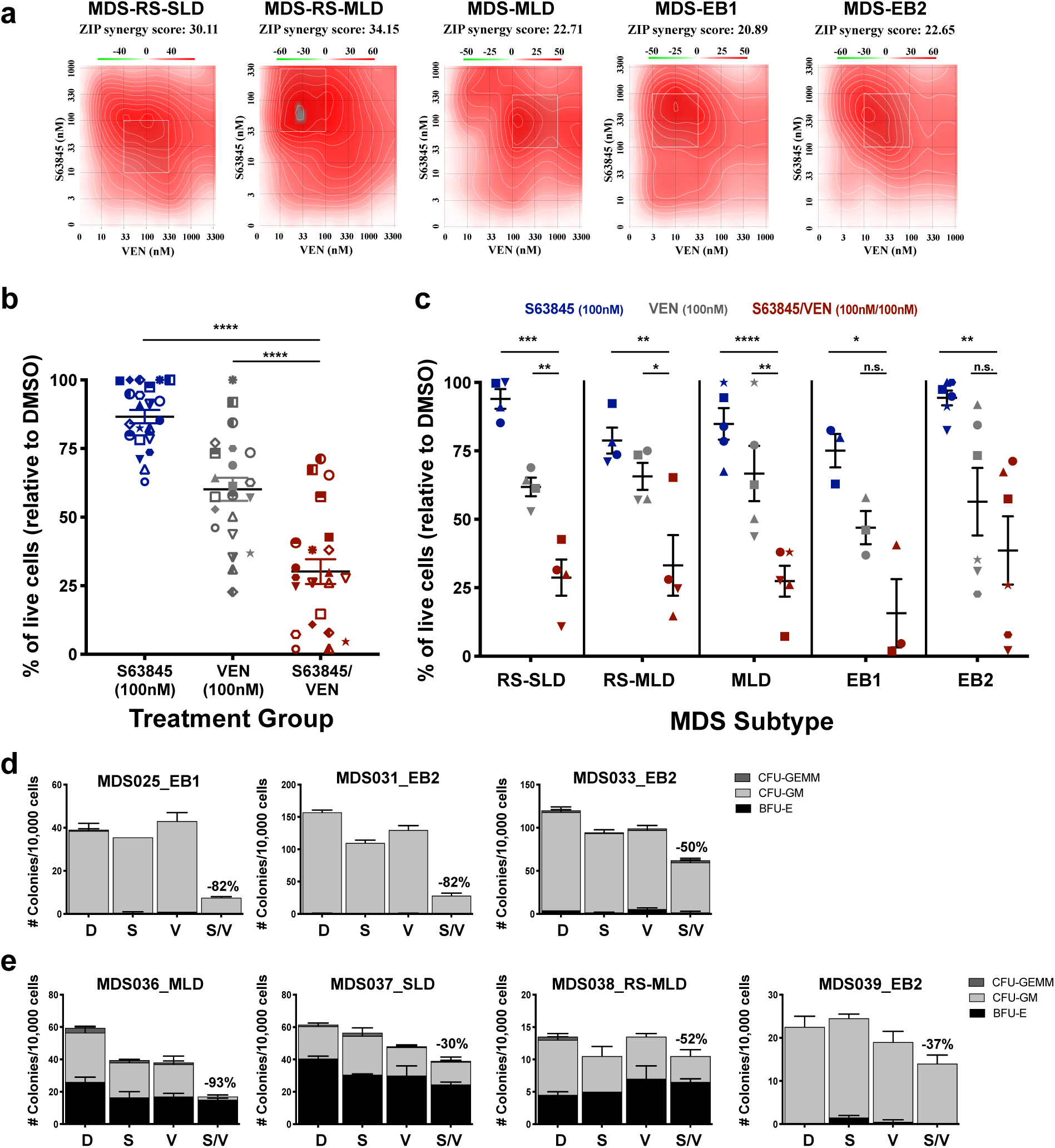
Dual inhibition of MCL1 and BCL2 is synergistic in MDS cells resulting in increased apoptosis and loss of clonogenicity. **a** Cell viability of primary MDS samples was measured by CellTiter-Glo at 48 hr after treatment with three-fold dilutions of S63845, venetoclax (VEN) or a combination of both. Contour plots of synergy scores generated from the cell viability dose matrix of S63845 and VEN using the zero interaction potency (ZIP) model. The synergy scores were represented by pseudocoloring 2-D contour plots over the dose matrix, giving rise to the overall synergy landscape. Red color indicates synergy, while green color indicates antagonism for the various concentrations of the S63845/VEN combination. (Note different pseudocoloring scale for ZIP synergy scores between the samples) **b-c** Apoptosis in CD34^+^ cells was measured by annexin V/PI staining using flow cytometry after 24 hr of treatment in all samples combined (**b**) or broken down into MDS subtypes (**c**). The ratio of live cells relative to DMSO control were calculated. Individual patient samples in b-c are represented by symbols with means ± SEM, and statistical comparisons between S63845 or VEN monotherapy and the combination are shown (two-tailed t-test; n.s. not significant, *p < 0.05, **p <0 .01, ***p<0 .001, ****p < 0.0001). **d-e** Colony forming assays of frozen (**d**) or freshly-obtained (**e**) MDS cells. The number of colonies per 1 × 10^4^ cells plated were calculated for each of the treated samples and for a DMSO control. The percent reduction of CFU-GM colonies in the combination treatment group compared to the DMSO control group is listed above each combination bar. Data is represented as an average of two replicates ± SEM. MDS-RS-SLD is MDS with ring sideroblast with single lineage dysplasia; MLD is multi-lineage dysplasia; EB1 is excess blasts 5-9%; EB2 is 10-19% blasts; CFU-GEMM is colony-forming unit – granulocyte, erythroid, macrophage, megakaryocyte; CFU-GM is colony-forming unit – granulocyte, macrophage; BFU-E is burst-forming unit – erythroid.

### Combined MCL1 and BCL2 inhibition reduces human MDS cell engraftment in MISTRG6 mice

We further validated the efficacy of S63845+VEN combination treatment using *in vivo* MDS patient-derived xenotransplants (PDX). We have previously shown successful establishment of MDS PDX with faithful clonal representation in cytokine humanized ‘MISTRG’ mice. MISTRG mice carry humanized alleles for <u>M</u>-CSF^h/h^, IL-3/GM-CSF^h/h^, <u>S</u>IRPα^h/m^, and <u>T</u>PO^h/h^, on the <u>R</u>AG2^-/-^ <u>γ</u>c^-/-^ background and are viable, healthy and fertile. Gene-humanizations via knock-in significantly improve the engraftment, differentiation and maintenance of human cells from MDS and other hematologic malignancies^20-22^. As MDS is an inflammatory disorder with elevated IL-6 levels and is highly dependent on its microenvironment^23,24^, we made use of a next-generation version of MISTRG with humanization via targeted insertion of human interleukin-6 (*IL-6*), in short, ‘MISTRG6’. MISTRG6 have been shown to efficiently engraft healthy human huCD34^+^ and malignant cells^21,25^. Before testing the effects of combined BCL2 and MCL1 inhibition experiments in MDS PDXs, we performed toxicity evaluations in MISTRG6 mice. Mice were treated with vehicle, VEN (15 mg/kg; 5 days on, 2 days off), or a combination of VEN (15 mg/kg; 5 days on, 2 days off) and S63845 (12.5 mg/kg; 2 days on, 5 days off) for a total of 2 weeks. Measurements were taken prior to starting treatment (Pre), 1 week, and 2 weeks after treatment to determine weight, red blood cell count, hemoglobin in complete blood count, white blood cell count, and platelet count. No detrimental effects were seen in the mouse hematopoietic cells in MISTRG6 mice (Supplementary Figure 2). Next, we confirmed engraftment of selected MDS patient samples from Table 1 that had sufficient cells to perform *in vivo* studies. All 3 samples engrafted in the MISTRG6 mice, at varying levels in the bone marrow (BM) (Fig. 2a-b), and were largely comprised of the myeloid lineage (Fig. 2c). To maximize the information we could obtain with this limited resource, we used these test engraftment mice to perform a pilot study comparing vehicle treatment to combination BCL2+MCL1 inhibition. This preliminary experiment hinted that combination treatment would decrease MDS engraftment in the BM (Supplementary Figure 3).

**Fig. 2.**
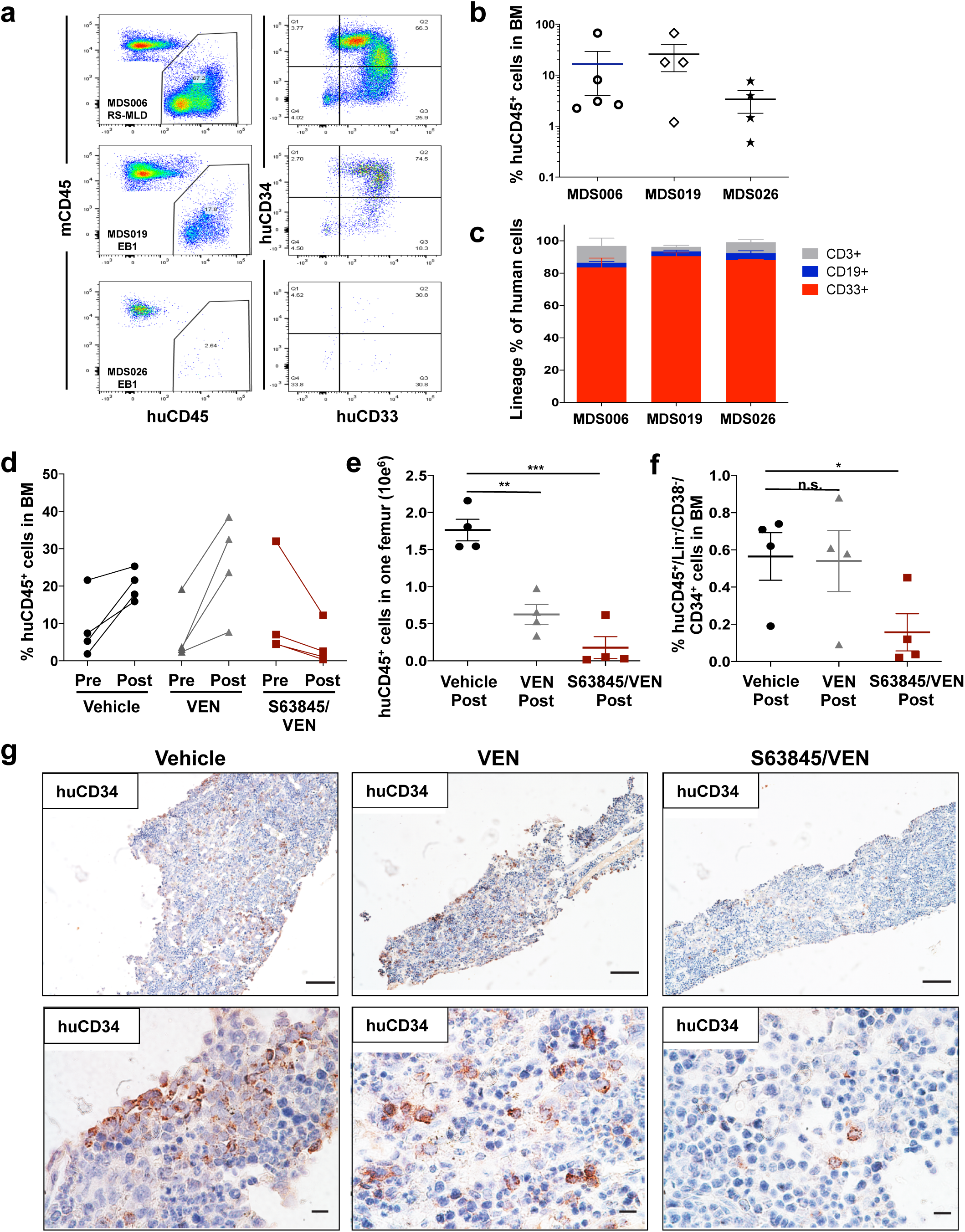
Evidence of reduced MDS engraftment in MISTRG6 mice after co-inhibition of BCL2 and MCL1.**a-c** Confirmed engraftment of 3 primary MDS patient cells in MISTRG6 mice. **a** Flow cytometric analysis of mouse vs human CD45 (mCD45 vs. huCD45; left panels) and huCD34 vs huCD33 (right panels). **b** Percent of huCD45^+^ cells in the bone marrow (BM) for each patient sample. **c** Relative distribution of myeloid CD33^+^ (red), B-lymphoid CD19^+^ (blue), and T-lymphoid CD3^+^ (gray) cells as percent of human CD45^+^ cells for each patient sample. **d-g** Patient sample MDS019-EB1 was allowed to engraft in MISTRG6 mice for 12 weeks before beginning treating with vehicle, VEN (15 mg/kg; 5 days on, 2 days off), or a combination of VEN (15 mg/kg; 5 days on, 2 days off) and S63845 (12.5 mg/kg; 2 days on, 5 days off) for a total of 4 weeks (n = 4 per treatment group). The percent (**d**) or total number (**e**) of huCD45^+^ cells in the BM were compared between pre and post-treatment for each treatment group. **f** After completion of the treatment, the percent of huCD45^+^/lin^-^/huCD38^-^/huCD34^+^ in the BM was determined. **g** Representative histologic images of BM from mice in d-f that were treated with vehicle, VEN, or S63845/VEN and underwent immunohistochemical staining for huCD34 (top panel, original magnification 10x, scale bars: 100μm; lower panel, original magnification 60x, scale bars:10μm). The data in **e-f** are represented by means ± SEM, and statistical comparisons between vehicle and VEN or combination treatments are shown (two-tailed t-test; n.s. not significant, *p < 0.05, **p <0 .01, ***p<0 .001).

Since MDS019 led to the most consistent engraftment (Fig. 2a-b), remaining cells from this patient sample (MDS-EB1 subtype) were allowed to engraft in MISTRG6 mice for 16 weeks before beginning treatment with vehicle, VEN (15 mg/kg; 5 days on, 2 days off), or a combination of VEN (15 mg/kg; 5 days on, 2 days off) and S63845 (12.5 mg/kg; 2 days on, 5 days off) for a total of 4 weeks (n = 4 per treatment group). The percent or total number of human CD45^+^ (huCD45^+^) cells in the BM were compared between pre- and post-treatment for each treatment group (Fig. 2d-e, respectively). Both of these analyses demonstrated a marked reduction in huCD45^+^ cells in the BM after combined BCL2+MCL1 inhibition, with almost a complete loss of huCD45^+^ cells. Furthermore, after the completion of the treatment, measurement of the percent of huCD45^+^/lin^-^/huCD38^-^/huCD34^+^ HSPCs in the BM (Fig. 2f), indicated this combined treatment was successful at specifically reducing the HSPCs that are known to drive the propagation of MDS, further exemplified by reduced CD34^+^ staining by immunohistochemistry in the bone marrow (Fig. 2g).

## Discussion

Study of MDS HSPCs *in vitro* is challenging. For the most part, MDS cell lines are contrived and limited, and patient samples are required^26-28^. *Ex vivo* study of MDS patient samples is also limited by sample heterogeneity, availability, and the practical consideration of how few cells are available with the background of marrow failure. Nonetheless, we have performed a comprehensive analysis on a cohort of MDS patient samples to investigate the value of targeting anti-apoptotic proteins for the treatment of MDS. Overall, our data validated previous findings that higher blast count MDS subtypes (EB1 and EB2) are more sensitive to VEN monotherapy than low blast count subtypes^1,4^ and that mutations associated with VEN resistance in AML may be similarly relevant in MDS. We also discovered, importantly, that MCL1 inhibtion may be effective for all MDS subtypes. Moreover, drug synergy from combining BCL2 and MCL1 inhibitors can be achieved across all subtypes and mutational profiles of MDS in this model, even in the presence of *RAS* family mutations that are thought to confer resistance to VEN monotherapy (eg. *CBL* and *PTPN11* in this cohort)^14-18^.

The *in vivo* reduction of MDS progenitor in the BM from MDS patient-derived xenografts is the first successful test of selective antiapoptotic inhibitors in an *in vivo* MDS PDX model. Prior patient-derived MDS xenograft models have demonstrated inefficient engrafment potential in multiple murine backgrounds and often do not recapitulate the disease characteristics^27,29-37^. Other studies have shown mixed results when attempting to improve MDS cell engraftment into mice with co-injection of normal or MDS patient-derived mesenchymal stromal cells^35,37^. The MISTRG mice overcame the obstacles presented by other models by expressing human cytokines in place of their murine counterparts at the endogenous locus to provide physiological expression of these human cytokines and MISTRG6 mice further provide the otherwise non-crossreactive critical inflammatory cytokine, IL-6, which is expressed in mesenchymal stromal cells in MDS and other hematologic malignancies^20^. Building on this model allowed us to successfully engraft MDS patient cells to test the effects of anti-apoptotic inhibitors *in vivo*. Taken together, these MDS PDX experiments inform on the selective utility of BCL2 inhibitor monotherapy in MDS and provide promise for the use of combined BCL2+MCL1 inhibition for the treatment of all MDS subtypes.

BCL2 inhibition is changing the standard of care in AML; refining the design of clinical trials testing BCL2 and MCL1 inhibitors in MDS and the precision of patient selection for therapy is a great priority. While combination of BCL2 and MCL1 inhibition may have a dose-dependent impact on normal hematopoiesis, experimental evidence seems to indicate that toxicity on normal CD34^+^ cells *in vitro*^38^ and *in vivo*^11^ is limited at clinically meaningful doses. The data presented here suggest dual BCL2+MCL1 inhibition may be considered for the treatment of all MDS subtypes regardless of mutational status, even those with mutations that may be less likely to respond to VEN monotherapy (eg. *TP53* or *RAS* family mutations)^14-18^, and carefully designed clinical trials with flexible dosing schedules amenable to MDS are warranted.

## Methods

### Patient samples

Experiments were conducted on primary MDS patient samples accessed from the Vanderbilt-Ingram Cancer Center Hematologic Malignancy Tumor Bank in accordance with the tenets of the Declaration of Helsinki and approved by the Vanderbilt University Medical Center Institutional Review Board.

### In vitro growth inhibition assays and determination of GI_50_ values

Cells were plated into a 384-well dish with IMDM + 10% FBS + 1% Pennicillin/Streptomycin + cytokines and supplements (10ng/ml hIL3, 10ng/ml hFLT3 ligand, 10ng/ml hTPO, 10ng/ml hSCF, 5ng/ml hIL6, 4μg/ml hLDL, 10μM 2-Mercaptoethanol, and 1x L-GlutaMAX) and treated for 48 hours with an MCL1 inhibitor (S63845; Chemietek, Indianapolis, IN), BCL2 inhibitor (venetoclax; Chemietek), or BCL-xL inhibitor (A-1155463; Chemietek) at concentrations from 3nM to 10µM. For combination studies, 3-fold dilution matrices of each agent were used between 10nM and 3.3µM. Cell viability was measured using CellTiterGlo (Promega, Madison, WI), and the relative luminescence units (RLU) were measured with a micro-plate reader (BioTek, Winooski, VT). Viability was defined as the percent of RLU of each well compared to the RLU of cells treated with DMSO vehicle. The 50% growth inhibition concentration (GI_50_) values were determined using linear regression of double-log transformed data.

### Drug combination calculation of synergy

The effects of S63845+VEN were calculated using the Zero Interaction Potency (ZIP) model, which compares observed and expected combination effects^19,39^.

### Assessment of apoptosis

Apoptosis was analyzed 24 hours after treatment (same concentrations as the CFU assay) using a FITC-AnnexinV/Propidium Iodide (PI) Apoptosis Detection kit per manufacter’s protocol (BD Biosciences, San Jose, CA) and as previously described using the anti-human CD34 antibody11 (Clone 581; Biolegend, San Diego, CA).

### Colony-forming Unit (CFU) assays

Fresh MDS cells were treated with 100nM VEN, 100nM S63845, or 100nM of each compound for S63845+VEN treatments. After 24 hours of treatment, cells were plated in Methocult H4034 Optimum (Stem Cell Technologies, Vancover, Canada) at a density of 1-2 × 10^4^ cells per mL. Cryopreserved MDS cells were thawed and cultured in StemSpan™ + cytokines (10ng/ml hIL3, 10ng/ml hSCF, 5ng/ml hIL6) overnight before plating cells in Methocult H4034 Optimum (Stem Cell Technologies) at a density of 1-2 × 10^4^ cells per mL with DMSO, 100nM VEN, 100nM S63845, or 100nM of each compound for S63845+VEN treatments. Colonies were counted and identified after 12-14 days, and counts were normalized per 1 × 10^4^ for graphical representation. After colony counts were completed, the duplicate plates for each treatment group were washed with PBS + 0.5% bovine serum albumin and pooled to obtain live cell number counts via trypan blue exclusion from the methocult assays.

### Generation and analysis of MDS PDX

All animal experiments were approved by the Institutional Animal Care and Use Committee of Yale University. Mouse breeding: MIS^h/h^TRG6 mice with homozygous knockin replacement of the endogenous mouse Csf1, Il3, Csf2, Tpo, Il6 and Sirpa with their human counterparts were bred to MITRG mice to generate human cytokine homozygous and hSIRPA heterozygous mice (MIS^h/m^TRG6, labeled MISTRG6 throughout the study). Mice were maintained on continuous treatment with enrofloxacin in the drinking water (0.27□mg/mL, Baytril, Bayer Healthcare). MISTRG/MISTRG6 mice will be available via MTA and requests should be sent to.

Xenografting: Newborn MISTRG6 mice (3 days of age) were X-ray irradiated (X-RAD 320 irradiator) with 2□× □150□cGy 4□h apart. MDS patient BM CD34-selected cells were incubated with a murine anti-human CD3 antibody (clone Okt3, BioXCell, NH, USA) at 5□µg/100□µl for 10□min at room temperature prior to injection. Cells were injected intrahepatically in a volume of 20□µL with a 22-gauge Hamilton needle (Hamilton, Reno, NV). Engraftment levels were assessed via BM aspiration at 12/ 16 weeks post-transplantation to assign treatment groups.

Flow cytometric analysis: Engraftment of human CD45^+^ cells and their subsets were determined by flow cytometry. In brief, cells were isolated from engrafted mice, blocked with human/murine Fc block (BD Pharmingen, CA, US), and stained with combinations of antibodies all purchased from Biolegend: HSPC panel: APC/Cy7 mCD45 (30-F11, 1:300), APC/Cy7 mTer119 (Ter-119, 1:300), BV510 hCD45 (HI30, 1:100), BV421 huCD38 (HIT2, 1:100), PE huCD34 (561, 1:100), PE/Cy7 huCD10 (HI10a, 1:100); human engraftment panel: APC/Cy7 mCD45 (30-F11, 1:300), APC/Cy7 mTer119 (Ter-119, 1:300), BV510 hCD45 (HI30, 1:100), FITC huCD3 (OKT3, 1:100), PE/Cy7 huCD19 (HIB19, 1:100), APC huCD33 (WM53, 1:100), PE huCD34 (561, 1:100). Data were acquired with FACSDiva on an LSR Fortessa (BD Biosciences) equipped with 5 lasers and analyzed with FlowJo V10 software.

Histologic analysis: Tissues were fixed in BBC Biochemical B-PLUS FIX Fixative solution (Thermo Fisher Scientific, MA, USA) and embedded in paraffin. Femurs were decalcified with Formic Acid Bone Decalcifier (Decal Chemical, NY, USA). Paraffin blocks were sectioned at 4□μm and stained with hematoxylin and eosin (H&E) and antigen-specific antibodies by the Yale Clinical Pathology and Yale Pathology Tissue Services. Antibodies: human CD45, Leucocyte Common Antigen, clone PD7/26 + 2B11, Dako; human CD34 Class II, clone QBEnd 10, Dako. Images were acquired using Nikon Eclipse 80i microscope.

### Next generation sequencing (NGS) and analysis

NGS results available in clinical records were used when available. When clinical results were not available, NGS was performed on DNA extracted from bio-banked MDS samples using QIAamp DNA Blood Kit (Qiagen) and Trusight Myeloid Panel (Illumina, San Diego, CA) with the exact same genetic regions of interest as the clinically utilized panel, OnkoSight™ (Bio-reference Laboratories, Elmwood Park, NJ). Alignment to hg19 and variant calling was performed using Illumina Enrichment app in BaseSpace, with restriction to minimum depth of 500x and variant allele frequency of 5%, in accordance with the manufacturer’s parameters. Variant annotation was done with a modified version of vcf2maf.pl (https://github.com/mskcc/vcf2maf), a wrapper around ensembl’s Variant Effect Predictor. Multiple annotation packages, including dbNSFP, ExAC, COSMIC, dbSNP, ClinVar, SIFT, Polyphen2, GERP, and CADD were used. Variants designated “common”, following the protocol established by AACR GENIE^40^, were considered common polymorphisms.

### Statistical analysis

Data are shown as mean ± SEM and analyzed using Prism 8 (GraphPad Software, La Jolla, CA). Correlation analysis was determined using Spearman non-parametric correlation and statistical analysis was performed using an ANOVA (P<0.0001) followed by the student 2-tailed *t*-test. P values of < 0.05 were considered to be statistically significant (n.s. not significant, *p < 0.05, **p <0 .01, ***p<0 .001, ****p < 0.0001).

## Acknowledgements

This work was generously supported by the E.P. Evans Foundation, the Biff Rittenberg Foundation, and the Leukemia and Lymphoma Society for which, MRS is a Clinical Scholar. S.H was supported by the NIH/NIDDK (R01DK102792), the Frederick A. Deluca Foundation, the Edward P. Evans Foundation, and the Department of Defense. Y.S. was supported by a pilot grant from the Yale Cooperative Center of Excellence in Hematology (YCCEH; NIDDK U54DK106857) and the Young Scientists Fund of the National Natural Science Foundation of China (Grant No. 81800122). This study was in part supported by the Animal Modeling Core of the Yale Cooperative Center of Excellence in Hematology (NIDDK U54DK106857).

Patient samples were garnered from the Vanderbilt-Ingram Cancer Center (VICC) Hematologic Malignancy Tumor Bank. The Vanderbilt-Ingram Cancer Center grant (NCI P30 CA068485-19) supported this work. The REDCap database tool is supported by grant UL1 TR000445 from NCATS/NIH. Flow Cytometry experiments were performed in the VUMC Flow Cytometry Shared Resource. The VUMC Flow Cytometry Shared Resource is supported by the Vanderbilt Ingram Cancer Center (P30 CA68485) and the Vanderbilt Digestive Disease Research Center (DK058404). We thank Yale Pathology Tissue Services especially A. Brooks for research histology services. We thank Yale Animal Resources Center, especially P. Ranney, J. Fonck for animal care. We thank Yale Flow Cytometry Core, especially Lesley Divine, Diane Trotta and Chao Wang for flow cytometry analysis.

## Author Contributions

M.A.F., Y.S., and R.G. designed and performed experiments. M.P.A., and M.T.V. performed experiments. M.A.F., Y.S., R.G., M.P.A., M.T.V., M.A.C., B.N.S, T.P.S., S.H., and M.R.S. analyzed data. M.A.F. and M.R.S. performed statistical analysis and wrote the manuscript. S.H. and M.R.S. designed and supervised the study. All authors reviewed and edited drafts of the manuscript and approved the final version of the manuscript.

## References

1. Reidel V, Kauschinger J, Hauch RT, et al. Selective inhibition of BCL-2 is a promising target in patients with high-risk myelodysplastic syndromes and adverse mutational profile. Oncotarget. 2018;9(25):17270–17281.

2. Parker JE, Mufti GJ, Rasool F, Mijovic A, Devereux S, Pagliuca A. The role of apoptosis, proliferation, and the Bcl-2-related proteins in the myelodysplastic syndromes and acute myeloid leukemia secondary to MDS. Blood. 2000;96(12):3932–3938.

3. Invernizzi R, Pecci A, Bellotti L, Ascari E. Expression of p53, bcl-2 and ras oncoproteins and apoptosis levels in acute leukaemias and myelodysplastic syndromes. Leuk Lymphoma. 2001;42(3):481–489.

4. Jilg S, Reidel V, Muller-Thomas C, et al. Blockade of BCL-2 proteins efficiently induces apoptosis in progenitor cells of high-risk myelodysplastic syndromes patients. Leukemia. 2016;30(1):112–123.

5. Bhola PD, Letai A. Mitochondria-Judges and Executioners of Cell Death Sentences. Mol Cell. 2016;61(5):695–704.

6. Letai A, Bassik MC, Walensky LD, Sorcinelli MD, Weiler S, Korsmeyer SJ. Distinct BH3 domains either sensitize or activate mitochondrial apoptosis, serving as prototype cancer therapeutics. Cancer Cell. 2002;2(3):183–192.

7. Hanahan D, Weinberg RA. Hallmarks of cancer: the next generation. Cell. 2011;144(5):646–674.

8. DiNardo CD, Pratz K, Pullarkat V, et al. Venetoclax combined with decitabine or azacitidine in treatment-naive, elderly patients with acute myeloid leukemia. Blood. 2019;133(1):7–17.

9. Wei AH, Strickland SA, Hou J-Z, et al. Venetoclax combined with low-dose venetoclax for previously untreated patients with acute myeloid leukemia: Results from a phase Ib/II study. Journal of Clinical Oncology. 2019;In press.

10. Pan R, Ruvolo VR, Wei J, et al. Inhibition of Mcl-1 with the pan-Bcl-2 family inhibitor (-)BI97D6 overcomes ABT-737 resistance in acute myeloid leukemia. Blood. 2015;126(3):363–372.

11. Ramsey HE, Fischer MA, Lee T, et al. A Novel MCL1 Inhibitor Combined with Venetoclax Rescues Venetoclax-Resistant Acute Myelogenous Leukemia. Cancer Discov. 2018;8(12):1566–1581.

12. Konopleva M, Pollyea DA, Potluri J, et al. Efficacy and Biological Correlates of Response in a Phase II Study of Venetoclax Monotherapy in Patients with Acute Myelogenous Leukemia. Cancer Discov. 2016;6(10):1106–1117.

13. Malcovati L, Papaemmanuil E, Ambaglio I, et al. Driver somatic mutations identify distinct disease entities within myeloid neoplasms with myelodysplasia. Blood. 2014;124(9):1513–1521.

14. Pei S, Pollyea DA, Gustafson A, et al. Monocytic Subclones Confer Resistance to Venetoclax-Based Therapy in Patients with Acute Myeloid Leukemia. Cancer Discov. 2020;10(4):536–551.

15. Chyla B, Daver N, Doyle K, et al. Genetic Biomarkers Of Sensitivity and Resistance to Venetoclax Monotherapy in Patients With Relapsed Acute Myeloid Leukemia. Am J Hematol. 2018;93(8):E202–E205.

16. DiNardo CD, Tiong IS, Quaglieri A, et al. Molecular patterns of response and treatment failure after frontline venetoclax combinations in older patients with AML. Blood. 2020;135(11):791–803.

17. Chen X, Glytsou C, Zhou H, et al. Targeting Mitochondrial Structure Sensitizes Acute Myeloid Leukemia to Venetoclax Treatment. Cancer Discov. 2019;9(7):890–909.

18. Nechiporuk T, Kurtz SE, Nikolova O, et al. The TP53 Apoptotic Network Is a Primary Mediator of Resistance to BCL2 Inhibition in AML Cells. Cancer Discov. 2019;9(7):910–925.

19. Yadav B, Wennerberg K, Aittokallio T, Tang J. Searching for Drug Synergy in Complex Dose-Response Landscapes Using an Interaction Potency Model. Comput Struct Biotechnol J. 2015;13:504–513.

20. Song Y, Rongvaux A, Taylor A, et al. A highly efficient and faithful MDS patient-derived xenotransplantation model for pre-clinical studies. Nat Commun. 2019;10(1):366.

21. Das R, Strowig T, Verma R, et al. Microenvironment-dependent growth of preneoplastic and malignant plasma cells in humanized mice. Nat Med. 2016;22(11):1351–1357.

22. Saito Y, Ellegast JM, Rafiei A, et al. Peripheral blood CD34(+) cells efficiently engraft human cytokine knock-in mice. Blood. 2016;128(14):1829–1833.

23. Herold M, Schmalzl F, Zwierzina H. Increased serum interleukin 6 levels in patients with myelodysplastic syndromes. Leuk Res. 1992;16(6-7):585–588.

24. Boada M, Echarte L, Guillermo C, Diaz L, Touriño C, Grille S. 5-Azacytidine restores interleukin 6-increased production in mesenchymal stromal cells from myelodysplastic patients. Hematology, Transfusion and Cell Therapy. 2020.

25. Yu H, Borsotti C, Schickel JN, et al. A novel humanized mouse model with significant improvement of class-switched, antigen-specific antibody production. Blood. 2017;129(8):959–969.

26. Drexler HG, Dirks WG, Macleod RA. Many are called MDS cell lines: one is chosen. Leuk Res. 2009;33(8):1011–1016.

27. Rhyasen GW, Wunderlich M, Tohyama K, Garcia-Manero G, Mulloy JC, Starczynowski DT. An MDS xenograft model utilizing a patient-derived cell line. Leukemia. 2014;28(5):1142–1145.

28. Tohyama K, Tsutani H, Ueda T, Nakamura T, Yoshida Y. Establishment and characterization of a novel myeloid cell line from the bone marrow of a patient with the myelodysplastic syndrome. Br J Haematol. 1994;87(2):235–242.

29. Muguruma Y, Matsushita H, Yahata T, et al. Establishment of a xenograft model of human myelodysplastic syndromes. Haematologica. 2011;96(4):543–551.

30. Martin MG, Welch JS, Uy GL, et al. Limited engraftment of low-risk myelodysplastic syndrome cells in NOD/SCID gamma-C chain knockout mice. Leukemia. 2010;24(9):1662–1664.

31. Thanopoulou E, Cashman J, Kakagianne T, Eaves A, Zoumbos N, Eaves C. Engraftment of NOD/SCID-beta2 microglobulin null mice with multilineage neoplastic cells from patients with myelodysplastic syndrome. Blood. 2004;103(11):4285–4293.

32. Benito AI, Bryant E, Loken MR, et al. NOD/SCID mice transplanted with marrow from patients with myelodysplastic syndrome (MDS) show long-term propagation of normal but not clonal human precursors. Leuk Res. 2003;27(5):425–436.

33. Wunderlich M, Chou FS, Link KA, et al. AML xenograft efficiency is significantly improved in NOD/SCID-IL2RG mice constitutively expressing human SCF, GM-CSF and IL-3. Leukemia. 2010;24(10):1785–1788.

34. Krevvata M, Shan X, Zhou C, et al. Cytokines increase engraftment of human acute myeloid leukemia cells in immunocompromised mice but not engraftment of human myelodysplastic syndrome cells. Haematologica. 2018;103(6):959–971.

35. Medyouf H, Mossner M, Jann JC, et al. Myelodysplastic cells in patients reprogram mesenchymal stromal cells to establish a transplantable stem cell niche disease unit. Cell Stem Cell. 2014;14(6):824–837.

36. Meunier M, Dussiau C, Mauz N, et al. Molecular dissection of engraftment in a xenograft model of myelodysplastic syndromes. Oncotarget. 2018;9(19):14993–15000.

37. Rouault-Pierre K, Mian SA, Goulard M, et al. Preclinical modeling of myelodysplastic syndromes. Leukemia. 2017;31(12):2702–2708.

38. Moujalled DM, Pomilio G, Ghiurau C, et al. Combining BH3-mimetics to target both BCL-2 and MCL1 has potent activity in pre-clinical models of acute myeloid leukemia. Leukemia. 2018.

39. Ianevski A, He L, Aittokallio T, Tang J. SynergyFinder: a web application for analyzing drug combination dose-response matrix data. Bioinformatics. 2017;33(15):2413–2415.

40. Consortium APG. AACR Project GENIE: Powering Precision Medicine through an International Consortium. Cancer Discov. 2017;7(8):818–831.

