## Supplementary Figures for "MCL1 dependence across MDS subtypes and dual inhibition of MCL1 and BCL2 in MISTRG6 mice"

### Supplementary Figure 1

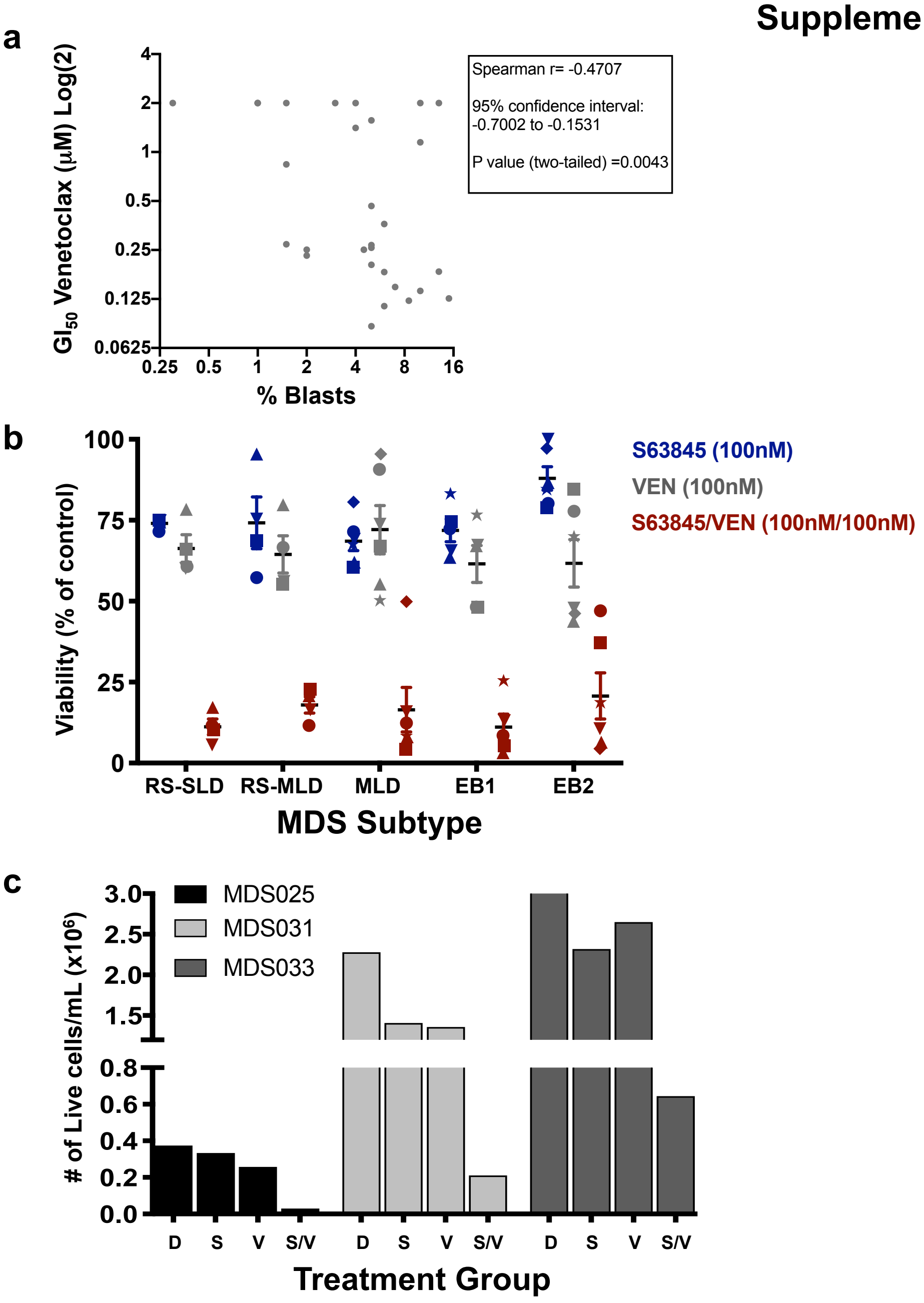


**Supplementary Figure 1.** Venetoclax sensitivity is correlated with blast percentage in MDS patient samples, and dual inhibition of MCL1 and BCL2 results in significant loss in cell viability. **a** Correlation analysis of $GI_{50}$ with venetoclax and blast percent in MDS patient samples. Blast percent in the bone marrow by IHC recorded in the clinical record, when available, or blast percent counted on morphologic assessment of the aspirate were used. $GI_{50}$ values greater than 2mM were capped at 2mM for graphing purposes. Spearman correlation and p value is displayed. **b** Cell viability was measured by CellTiter-Glo at 48 hr after treatment with 100nM of S63845, 100nM VEN, or 100nM S63845 + 100nM VEN and normalized to DMSO control. **c** After colony forming units were counted, replicate plates were washed and the number of live cells/mL from the combined plates were determined via trypan blue exclusion for each sample tested.

**Supplementary Table 1.** Average ZIP δ synergy score (based on the entire range of doses tested) and maximum δ score (based on the specific concentrations that have the highest synergy for each sample) for each MDS patient sample tested. Red shading indicates synergistic ZIP scores >0, with darker shading indicating higher synergy.

| **Sample ID** | **MDS Subtype** | **Average δ** Score | **Maximum** δ Score |
|---|---|---|---|
| **MDS001** | RS-SLD | 29.01 | 42.99 |
| **MDS002** | RS-SLD | 34.15 | 49.49 |
| **MDS003** | RS-SLD | 30.11 | 48.96 |
| **MDS004** | RS-SLD | 22.22 | 32.63 |
| **MDS005** | RS-MLD | 23.17 | 32.84 |
| **MDS006** | RS-MLD | 25.71 | 40.02 |
| **MDS008** | RS-MLD | 14.77 | 21.27 |
| **MDS009** | RS-MLD | 20.2 | 29.21 |
| **MDS010** | MLD | 22.98 | 37.55 |
| **MDS011** | MLD | 20.81 | 41.71 |
| **MDS013** | MLD | 19.72 | 31.01 |
| **MDS015** | MLD | 20.18 | 27.82 |
| **MDS016** | MLD | 32.23 | 43.13 |
| **MDS019** | EB1 | 21.64 | 34.93 |
| **MDS025** | EB1 | 32.55 | 48.44 |
| **MDS026** | EB1 | 12.54 | 19.53 |
| **MDS027** | EB1 | 19.54 | 32.61 |
| **MDS028** | EB1 | 20.89 | 35.01 |
| **MDS029** | EB2 | 25.9 | 38.47 |
| **MDS031** | EB2 | 19.07 | 26.57 |
| **MDS032** | EB2 | 27.19 | 41.87 |
| **MDS033** | EB2 | 17.98 | 32.71 |
| **MDS034** | EB2 | 22.65 | 37.57 |
| **MDS035** | EB2 | 11.01 | 16.09 |

#### Supplementary Figure 2

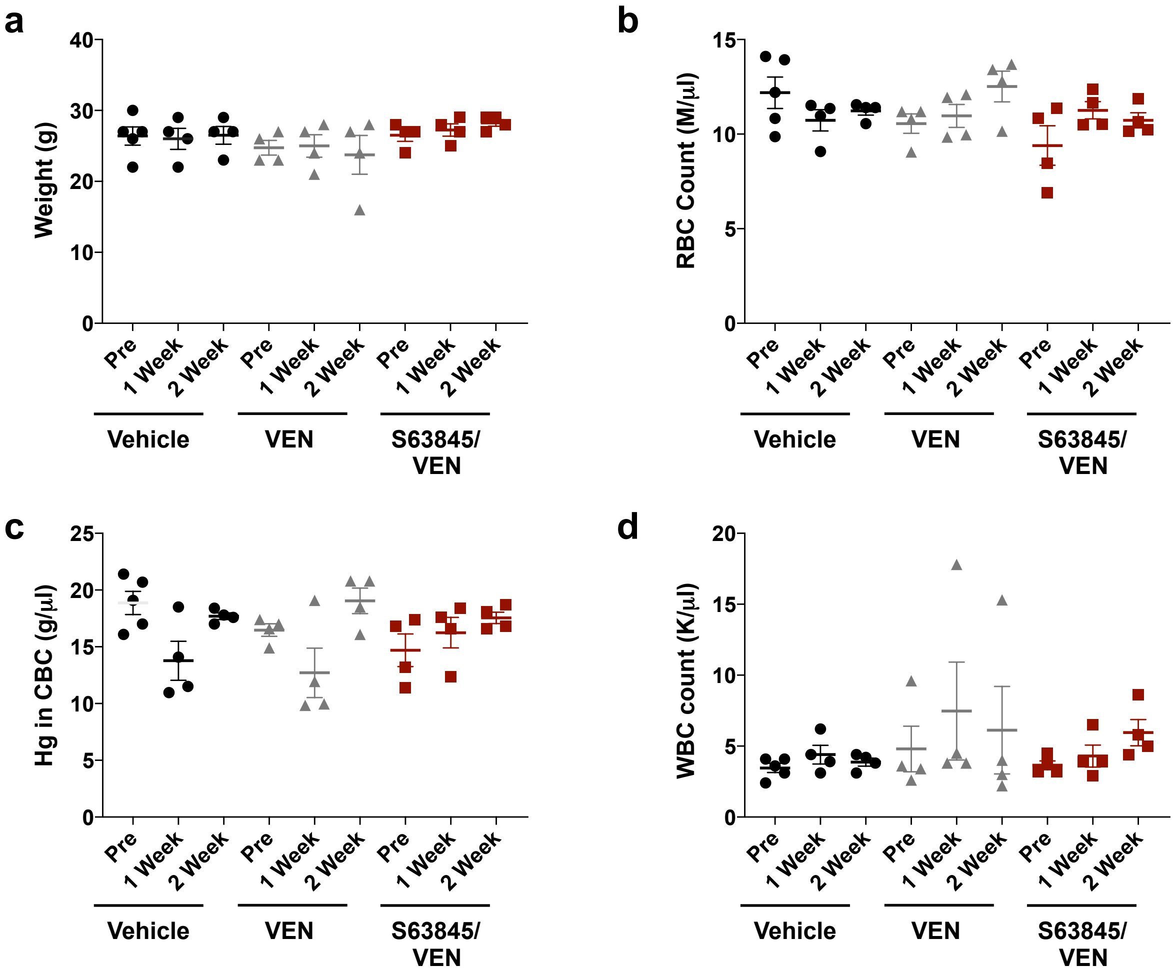


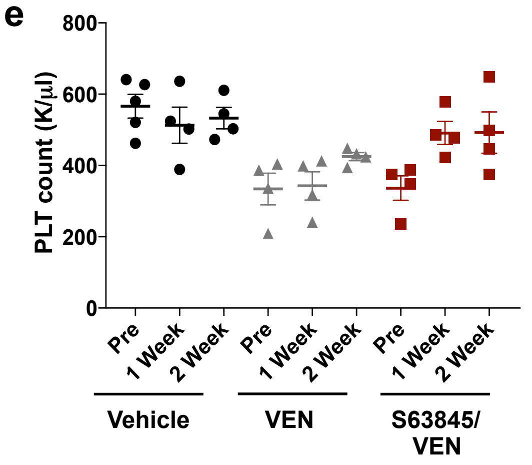


**Supplementary Figure 2.** MISTRG6 mice do not display hematopoietic toxicity when exposed to dual BCL2+MCL1 inhibition. **a-e** Toxicity experiments in the MISTRG6 (**M**-CSF$^{h/h}$, **I**L-3/ GM-CSF$^{h/h}$, h**S**IRPα$^{h/m}$, h**T**PO$^{h/h}$, and IL-**6** on the **R**AG2$^{-/-}$ **γ**c$^{-/-}$ background) immunocompromised animal model[20]. Mice were treated with vehicle, VEN (15 mg/kg; 5 days on, 2 days off), or a combination of VEN (15 mg/kg; 5 days on, 2 days off) and S63845 (12.5 mg/kg; 2 days on, 5 days off) for a total of 2 weeks. Measurements were taken prior to starting treatment (Pre), 1 week, and 2 weeks after treatment to determine weight (**a**), red blood cell (RBC) count (**b**), hemoglobin (Hg) in complete blood count (CBC) (**c**), white blood cell (WBC) count (**d**), and platelet (PLT) count (**e**).

#### Supplementary Figure 3

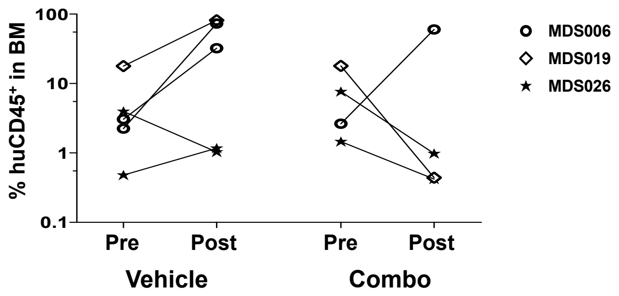


**Supplementary Figure 3.** Dual BCL2+MCL1 inhibition diminishes human MDS in bone marrow. **A**fter baseline engraftment was determined (pre), the mice from Fig. 2a-c were treated with vehicle, or a combination of VEN (15 mg/kg; 5 days on, 2 days off) and S63845 (12.5 mg/kg; 2 days on, 5 days off) for a total of 4 weeks before determining the percent of huCD45$^+$ engraftment (post) in the bone marrow (BM).
